# Novel, postnatal manifestation of an epidermal barrier defect in a mouse model of isolated sulfite oxidase deficiency

**DOI:** 10.1101/2024.03.14.584870

**Authors:** Lena Johannes, Matthias Rübsam, Julia Löhr, Xiaolei Ding, Sabine Eming, Carien M. Niessen, Günter Schwarz

## Abstract

Sulfite oxidase deficiency is a rare inborn error in metabolism leading to early childhood death due to rapidly progressing neurodegeneration. A new mouse model of sulfite oxidase deficiency carrying a homozygous deletion in the *Suox* gene resembles the human pathology in terms of neonatal death and elevation of sulfite and thiosulfate in plasma and urine, respectively. Homozygous *Suox*^*-/-*^ mice are initially born healthy, display growth retardation starting at postnatal day 4 and die in average at day 9.6. Here we report that *Suox*^*-/-*^ mice develop dry and scaly skin early postnatally, showing that sulfite oxidase is essential to maintain a functional skin barrier after birth. At postnatal day 5 *Suox*^*-/-*^ mice develop altered epidermal morphology and dysregulated early and late keratinocyte differentiation accompanied by increased stress response. We propose a sulfite-induced cleavage of disulfide bonds in key epidermal proteins essential for a functional barrier.

## To the Editor

Isolated sulfite oxidase deficiency (SOXD) is a rare metabolic disorder typically presenting with postnatal intractable seizures and progressive neurodegeneration, resulting in infant death if left untreated (Johnson & Duran, 2001). It is caused by a loss of function of the enzyme sulfite oxidase (SOX), which catalyzes the clearance of sulfite via oxidation to sulfate as the final step of cysteine catabolism (Irreverre et al., 1967). A loss in SOX activity results in an accumulation of toxic sulfite and elevated levels of the sulfur metabolites taurine, thiosulfate and S-sulfocysteine. Due to its structural resemblance to glutamate, S-sulfocysteine acts as an NMDA receptor agonist, thus causing vast excitotoxicity and manifestation of a progressive neurological phenotype (Kumar et al., 2017).

Recently, a mouse model for SOXD has been successfully established through inactivation of the *Suox* gene (Fu et al. 2024). Heterozygous animals develop without visible phenotype. Homozygous *Suox*^*-/-*^ mice are initially born healthy, but around postnatal day 4 start to display growth retardation and decreased neuromotor abilities, and die between P6 and P12 with an average survival time of 9.6 days. Interestingly, brain morphological analysis shortly before death did not reveal significant abnormalities, thus excluding neurodegeneration as the major cause of death, and indicating additional pathomechanisms and causes of death for *Suox*^*-/-*^ mice and human SOXD patients (Fu et al. 2024). Here we report that *Suox*^*-/-*^ mice develop dry and scaly skin early postnatally, showing that SOX is essential to maintain a functional skin barrier after birth.

Newborn (P0) *Suox*^*-/-*^ mice initially displayed normal skin, both macroscopically as well as in H&E stained histological sections of P0 back skin with the epidermis appearing similar to control mice (Fig. 1A, E). In agreement, immunofluorescence analysis revealed no change in localization of keratin (K)14 (Fig. 1B, F), marking the proliferating basal layer, and K10 (Fig.1C,G), a differentiation marker for all suprabasal layers (Fuchs & Green, 1980). Moreover, staining for proliferation marker Ki67 demonstrated similar proliferation between control and *Suox*^*-/-*^ newborn mice (Fig. 1D, H). Together, these data indicate that the skin develops normally *in utero* upon loss of SOX enzyme.

**Figure 1:**
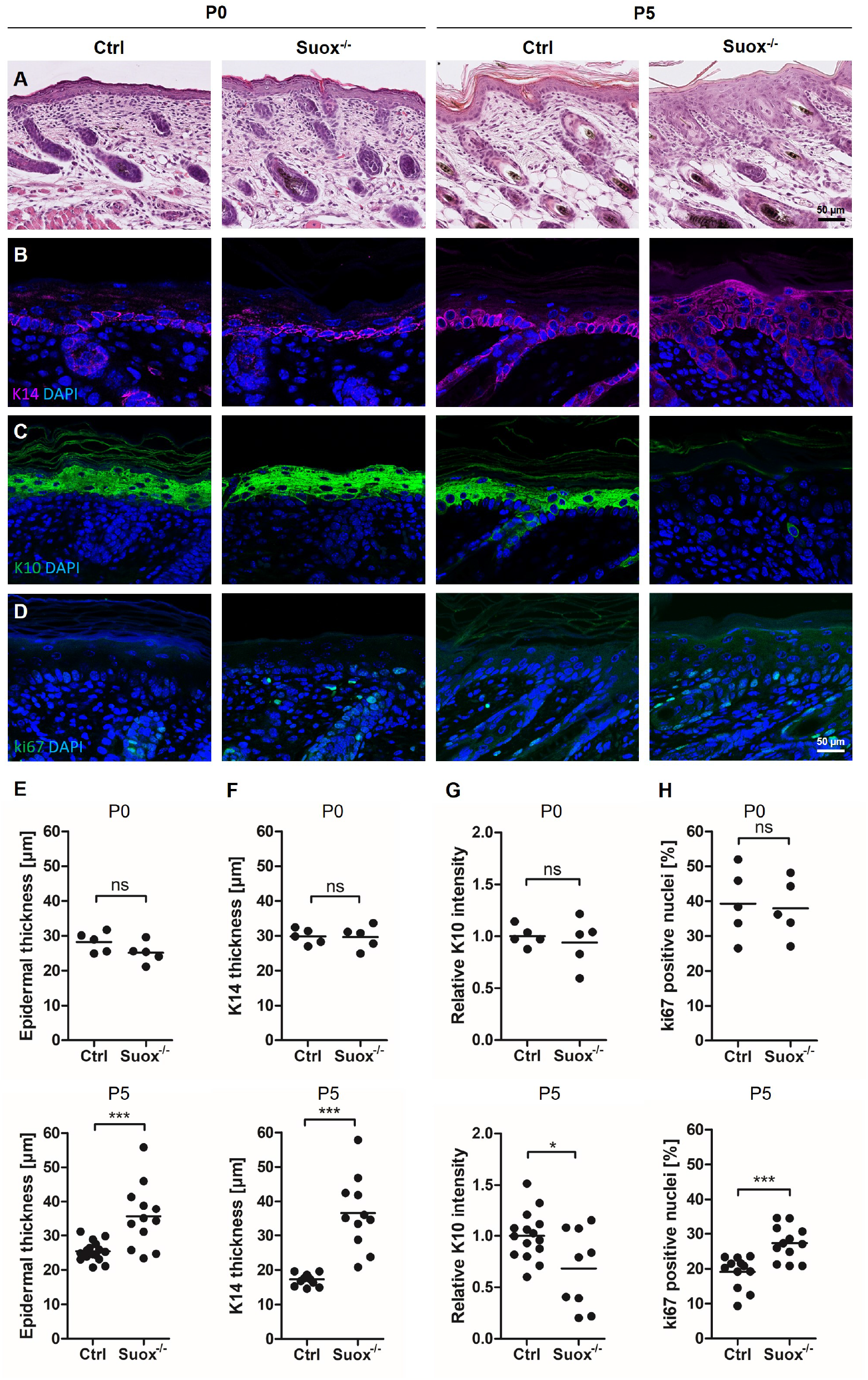
(previous page): *Suox*^*-/-*^ mice develop altered epidermal morphology and dysregulated early keratinocyte differentiation within the first days of life. (A) H&E stainings of back skin sections of *Suox*^*-/-*^ pups from P0 (left) and P5 (right). (B-D) Immunofluorescent stainings of paraffin-embedded back skin sections for (B) basal cell marker Keratin14 (K14), (C) early differentiation marker Keratin10 (K10) and (D) proliferation marker Ki67, each at P0 (left) and P5 (right). (E) Quantification of epidermal thickness (excluding SC). (F) Quantification of K14 thickness. (G) Quantification of K10 intensity. (H) Quantification of ki67-positive cells. Each data point represents one animal, per animal ten microscopic pictures were analyzed, \*\*\**P*<0.001, *\*P*<0.05 with Student’s *t*-test.

At P5, H&E staining revealed a significant increase in epidermal thickness in homozygous *Suox*^*-/-*^ animals (Fig. 1A, E). Closer examination indicated that this increase in thickness likely originates from a combination of enlarged cells in the *stratum basale* (SB) as well as an increase in cell number and layers of the suprabasal spinous layers (Fig. 1A). The *stratum granulosum* (SG) appeared thinner, with a reduced number of granules, while cells in the *stratum corneum* (SC) showed decreased structural organization and parakeratosis (Fig. 1A), suggesting changes in differentiation. In agreement, immunofluorescence analysis revealed a strong increase in suprabasal K14 staining (Fig. 1B, F) and a strong reduction in the differentiation marker K10 in suprabasal layers (Fig. 1C, G), indicating that cells that moved suprabasally failed to properly differentiate and retain basal identity. In agreement, the proliferation marker Ki67 was increased and no longer confined to the basal layer (Fig. 1D, H).

To directly confirm that differentiation was altered postnatally, we stained for late differentiation markers involucrin, loricrin and filaggrin. Whereas staining for loricrin was almost absent in P5 *Suox*^*-/-*^ skin (Fig. 2A), and for filaggrin largely reduced (Fig. 2B), involucrin was still similar to control mice (Fig. 2C). Furthermore, K6, a stress-induced keratin (Eckhardt et al., 2019), was increased in the intrafollicular epidermis (Fig. 2D), providing additional evidence for disturbed differentiation and the presence of stressed keratinocytes. Together, these data indicate that epidermal loss of SOX postnatally promotes proliferation and impairs differentiation, resulting in a morphologically altered barrier.

**Figure 2:**
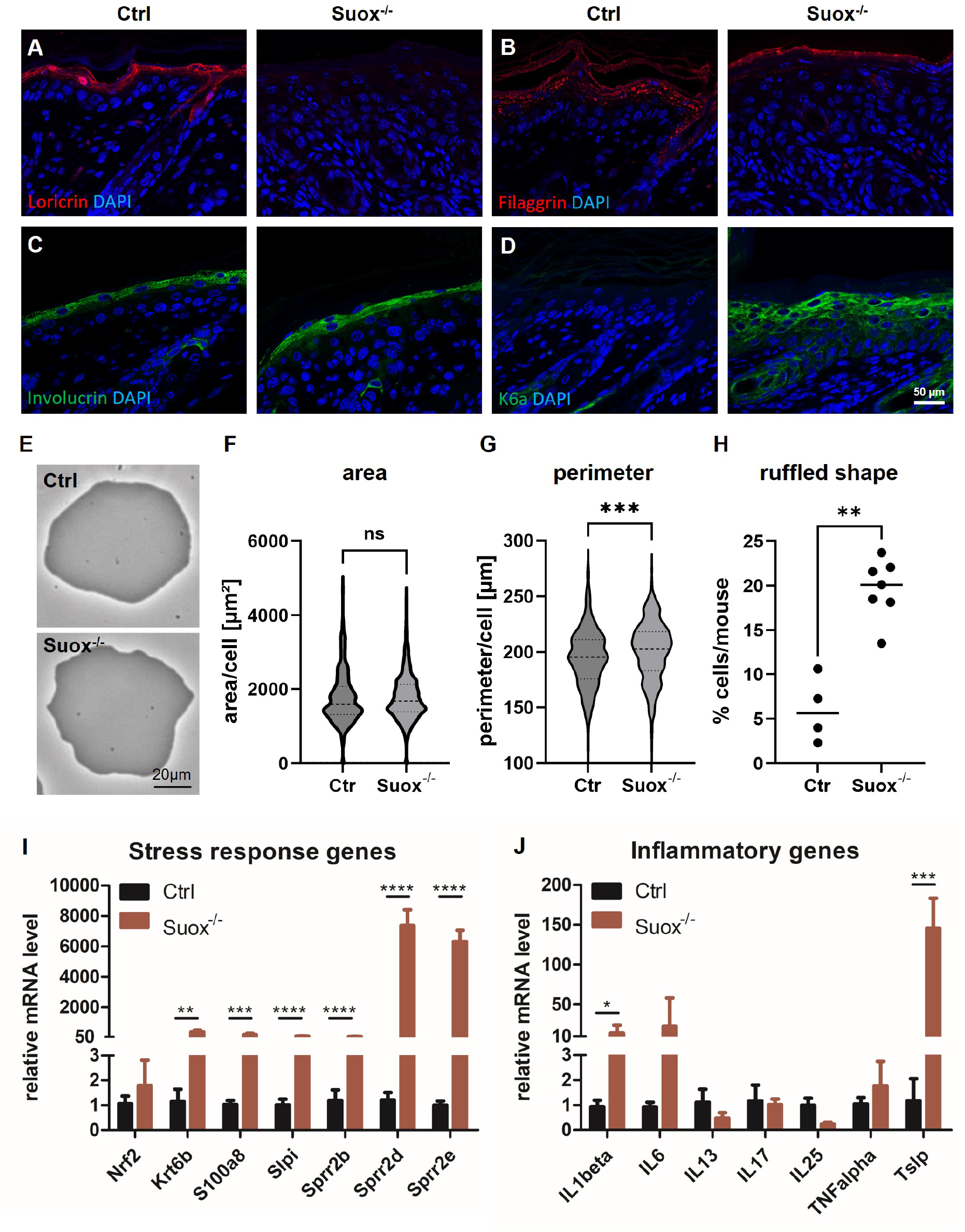
*Suox*^*-/-*^ mice develop dysregulated late keratinocyte differentiation and increased stress response within the first days of life. (A-D) Immunofluorescent stainings of paraffin-embedded back skin sections of control and *Suox*^*-/-*^ P5 for structural proteins and late differentiation markers (A) Loricrin, (B) Filaggrin, (C) Involucrin and (D) keratin K6a. (E) Isolated corneocyte appearance. (F) Quantification of corneocyte area, *P*=0.2258. (G) Quantification of corneocyte perimeter, *P*=0.0001. (F+G) >300 cells with Kolmogorov-Smirnov (accumulated from Ctr n=4, Suox^-/-^ n=7 biological replicates). (H) Quantification of ruffled corneocytes, \*\**P*=0.0061 with Mann–Whitney test; Ctr n=4, Suox^-/-^ n=7 biological replicates. (I+J) qRT-PCR analysis of stressed barrier (I) and inflammatory (J) genes on RNA isolated from the epidermis of control and *Suox*^*-/-*^ P5 mice. \*\*\**P*<0.001, \*\**P*<0.005, *\*P*<0.05 with Student’s *t*-test.

To investigate whether changes in epidermal terminal differentiation also altered barrier function upon loss of SOX, we isolated corneocytes from P5 mice and examined their appearance (Fig. 2E). Although average corneocyte area was still similar (Fig. 2F), the average perimeter was somewhat larger (Fig. 2G), likely due to an increase in the percentage of corneocytes with less smooth, more ruffled appearance (Fig. 2H), showing that the *stratum corneum* is abnormal and likely less functional. In agreement, real time PCR showed that next to K6 several markers (Slpi, S100a8, and several small proline-rich protein (Sprr) family members) known to be upregulated upon a dysfunctional barrier, were highly increased in *Suox*^*-/-*^ mice (Fig. 2I), further verifying the impairment of epidermal barrier integrity. Interestingly, *Sprr2* upregulation in loricrin^-/-^ mice is thought to compensate and maintain partial barrier function (Koch et al., 2000). Thus, the upregulation of several Sprrs observed here might equally function as an initial compensatory mechanism aiming to try and prevent the formation of a more severe epidermal phenotype.

An impaired barrier function often induces an innate inflammatory response. Although quantitative PCR analysis revealed no significant alteration in the expression of most inflammatory genes, *Il1b* mRNA levels were significantly elevated (Fig. 2J), whereas IL6 also showed a tendency for a strong increase. IL-1β was shown to induce the expression of K6a (Komine et al., 2001), while additionally activating transcription factor NF-κB, which can cause the downregulation of filaggrin and loricrin expression (Michaelidou et al., 2023). Moreover, mRNA levels of *Tslp*, a master regulator of type II skin immunity (Hasegawa et al. 2022), were also vastly elevated (Fig. 2J). In humans, cytokine TSLP was found to impair epidermal barrier integrity by upregulating the expression of nuclear cytokine IL-33, which in turn is involved in the downregulation of loricrin, filaggrin and K10 expression (Dai et al., 2021, 2022). Thus, the observed overexpression of IL-1β and TSLP presumably contributes to the altered expression patterns of respective marker proteins in our *Suox*^*-/-*^ mouse model as well.

Overall, our results suggest that late onset differentiation proteins such as the SPRR family together with hyperproliferation may act as a mechanism to counteract the postnatally occurring altered terminal differentiation and subsequent epidermal barrier dysfunction in *Suox*^*-/-*^ mice. Notably, our data revealed a variable phenotype, ranging from almost control-like to severely altered epidermal morphology at P5 (Fig. 1,2). As the life span of *Suox*^*-/-*^ mice typically lies between six and twelve days, the phenotypic severity at P5 may be linked to the age of onset and likely explains the variation in survival days. We therefore propose a progressive, postnatal loss of epidermal barrier integrity due to excessive accumulation of sulfite.

SOX shows moderate expression in the skin (Uhlén et al., 2015), but until this report, no role for its regulation in skin homeostasis was ascribed yet. However, sulfite, which accumulates upon SOX deficiency, can act as a nucleophile to cleave disulfide bonds (Kella & Kinsella, 1985). Disulfide bonds are crucial for epidermal barrier integrity, as these are important for the cross-linkage of keratin filaments and other structural proteins such as loricrin, the major component of the cornified cell envelope (Hohl et al., 1991; Suns & Green, 1978). In addition, a knock-in mouse model expressing K14 with a single cysteine-to-alanine mutation that reduced disulfide bond formation in the basal K14 protein, displayed changes in Yap-dependent transcription that impaired differentiation and barrier function, including suprabasal K14 expression and decreased K10, filaggrin and loricrin intensity (Guo et al., 2020). Thus, our results indicate that SOX postnatally regulates the extend of disulfide bonds in key epidermal proteins essential for a functional barrier. Further studies will be required to understand the contribution of cysteine modifications and unravel the exact pathomechanism.

## References

Dai, X., Muto, J., Shiraishi, K., Utsunomiya, R., Mori, H., Murakami, M., & Sayama, K. (2022). TSLP Impairs Epidermal Barrier Integrity by Stimulating the Formation of Nuclear IL-33/Phosphorylated STAT3 Complex in Human Keratinocytes. Journal of Investigative Dermatology, 142(8), 2100–2108.e5. 10.1016/J.JID.2022.01.005

Dai, X., Utsunomiya, R., Shiraishi, K., Mori, H., Muto, J., Murakami, M., & Sayama, K. (2021). Nuclear IL-33 Plays an Important Role in the Suppression of FLG, LOR, Keratin 1, and Keratin 10 by IL-4 and IL-13 in Human Keratinocytes. Journal of Investigative Dermatology, 141(11), 2646–2655.e6. 10.1016/J.JID.2021.04.002

Fu, C.-Y., Kohl, J.B., Liebsch, F., D’Andrea, D., Mai, M., Mellis, A.T., Kouroussis, E., Ditrói, T., Santamaria-Araujo, J.A., Yeo, S.Y., Endepols, H., Křížkov, M, Kožich, V., Barayeu, U., Akaike, T., Hennermann, J.B., Nagy, P., Filipovic, M., & Schwarz, G. (2024). Sulfite oxidase deficiency causes persulfidation loss and H2S release. BIORXIV/2024/584820

Fuchs, E., & Green, H. (1980). Changes in keratin gene expression during terminal differentiation of the keratinocyte. Cell, 19(4), 1033–1042. 10.1016/0092-8674(80)90094-X

Guo, Y., Redmond, C. J., Leacock, K. A., Brovkina, M. V, Ji, S., Jaskula-Ranga, V., & Coulombe, P. A. (2020). Keratin 14-dependent disulfides regulate epidermal homeostasis and barrier function via 14-3-3s and YAP1. 10.7554/eLife.53165

Hasegawa, T., Oka, T., Demehri, S (2022). Alarmin Cytokines as Central Regulators of Cutaneous Immunity. Front Immunol. 13:876515. doi:10.3389/fimmu.2022.876515. eCollection 2022.

Hohl, D., Mehrel, T., Lichti, U., Turner, M. L., Roop, D. R., & Steinert, P. M. (1991). Characterization of human loricrin. Structure and function of a new class of epidermal cell envelope proteins. Journal of Biological Chemistry, 266(10), 6626–6636. 10.1016/S0021-9258(18)38163-8

Irreverre, F., Mudd, S. H., Heizer, W. D., & Laster, L. (1967). Sulfite oxidase deficiency: Studies of a patient with mental retardation, dislocated ocular lenses, and abnormal urinary excretion of S-sulfo-l-cysteine, sulfite, and thiosulfate. Biochemical Medicine, 1(2), 187–217. 10.1016/0006-2944(67)90007-5

Johnson, J. L., & Duran, M. (2001). Molybdenum cofactor deficiency and isolated sulfite oxidase deficiency. In The Metabolic and Molecular Bases of Inherited Disease (pp. 3163–3177). Mcgraw-Hill Professional.

Kella, N. K. D., & Kinsella, J. E. (1985). A method for the controlled cleavage of disulfide bonds in proteins in the absence of denaturants. Journal of Biochemical and Biophysical Methods, 11(4–5), 251–263. 10.1016/0165-022X(85)90007-7

Koch, P. J., De Viragh, P. A., Scharer, E., Bundman, D., Longley, M. A., Bickenbach, J., Kawachi, Y., Suga, Y., Zhou, Z., Huber, M., Hohl, D., Kartasova, T., Jarnik, M., Steven, A. C., & Roop, D. R. (2000). Lessons from Loricrin-Deficient Mice: Compensatory Mechanisms Maintaining Skin Barrier Function in the Absence of a Major Cornified Envelope Protein. The Journal of Cell Biology, 151(2), 389. 10.1083/JCB.151.2.389

Kohl, J. B. (2019). A Novel Mouse Model of Sulfite Oxidase Deficiency: Pathological Changes in Cysteine and H2S Metabolism. In Department of Chemistry.

Komine, M., Rao, L. S., Freedberg, I. M., Simon, M., Milisavljevic, V., & Blumenberg, M. (2001). Interleukin-1 induces transcription of keratin K6 in human epidermal keratinocytes. Journal of Investigative Dermatology, 116(2), 330–338. 10.1046/j.1523-1747.2001.01249.x

Kumar, A., Dejanovic, B., Hetsch, F., Semtner, M., Fusca, D., Arjune, S., Santamaria-Araujo, J. A., Winkelmann, A., Ayton, S., Bush, A. I., Kloppenburg, P., Meier, J. C., Schwarz, G., & Belaidi, A. A. (2017). S-sulfocysteine/NMDA receptor-dependent signaling underlies neurodegeneration in molybdenum cofactor deficiency. J Clin Invest, 127(12), 4365–4378. 10.1172/JCI89885

Michaelidou, M., Redhu, D., Kumari, V., Babina, M., & Worm, M. (2023). IL-1α/β and IL-18 profiles and their impact on claudin-1, loricrin and filaggrin expression in patients with atopic dermatitis. Journal of the European Academy of Dermatology and Venereology. 10.1111/jdv.19153

Suns, T., & Green, H. (1978). Keratin filaments of cultured human epidermal cells. Formation of intermolecular disulfide bonds during terminal differentiation. 253(6), 2053–2060. 10.1016/S0021-9258(19)62353-7

Uhlén, M., Fagerberg, L., Hallström, B. M., Lindskog, C., Oksvold, P., Mardinoglu, A., Sivertsson, Å., Kampf, C., Sjöstedt, E., Asplund, A., Olsson, I. M., Edlund, K., Lundberg, E., Navani, S., Szigyarto, C. A. K., Odeberg, J., Djureinovic, D., Takanen, J. O., Hober, S., … Pontén, F. (2015). Tissue-based map of the human proteome. Science, 347(6220). 10.1126/SCIENCE.1260419/SUPPL_FILE/1260419_UHLEN.SM.PDF

